# Arousal and locomotion make distinct contributions to cortical activity patterns and visual encoding

**DOI:** 10.1101/010751

**Authors:** Martin Vinck, Renata Batista-Brito, Ulf Knoblich, Jessica A. Cardin

**Affiliations:** Yale University School of Medicine, Department of Neurobiology, New Haven, CT, USA; Kavli Institute of Neuroscience, Yale University, New Haven CT, USA

## Abstract

Spontaneous and sensory-evoked cortical activity is highly state-dependent, yet relatively little is known about transitions between distinct waking states. Patterns of activity in mouse V1 differ dramatically between quiescence and locomotion, but this difference could be explained by either motor feedback or a change in arousal levels. We recorded single cells and local field potentials from area V1 in mice head-fixed on a running wheel and monitored pupil diameter to assay arousal. Using naturally occurring and induced state transitions, we dissociated arousal and locomotion effects in V1. Arousal suppressed spontaneous firing and strongly altered the temporal patterning of population activity. Moreover, heightened arousal increased the signal-to-noise ratio of visual responses and reduced noise correlations. In contrast, increased firing in anticipation of and during movement was attributable to locomotion effects. Our findings suggest complementary roles of arousal and locomotion in promoting functional flexibility in cortical circuits.

****Abbreviations**:**

LFP: Local Field Potential
V1: primary visual cortex
PD: pupil diameter
FS: fast spiking
RS: regular spiking
FR: firing rate
L-on: locomotion onset
L-off: locomotion offset
L: locomotion
LE: early locomotion
LL: late locomotion
Q: quiescence
QE: early quiescence
QM: middle quiescence
QL: late quiescence
SNR: signal-to-noise ratio
ITI: Inter-trial-interval
LV: local coefficient of variation
ISI: inter-spike-interval
s.e.m.: standard error of the mean
SD: standard deviation
ID: Isolation Distance

## Introduction

Patterns of cortical activity differ dramatically across behavioral states, such as sleeping, anesthesia and waking^1–4^. Likewise, neural responses to sensory inputs depend strongly on ongoing patterns of internally generated activity^5–7^. The generation of multiple activity patterns associated with sleep and anesthesia states has been examined in great detail^2–4, 8–10^. However, relatively little is known about transitions between distinct waking states, such as quiescence, arousal, and focused attention.

Recent studies in rodents have contrasted inactive vs. active behavioral states, in particular quiescent vs. whisking^11–13^ or running^14–22^, and found profound differences in cortical activity patterns that resemble the effects of focused spatial attention in primates^23–27^. In mouse primary visual cortex (V1), locomotion is accompanied by altered firing rates, a reduction in low-frequency fluctuations in the membrane potential and local field potential (LFP), and an increase in LFP gamma-band oscillations^15–18, 22^. Enhanced firing rates during locomotion are particularly prominent in inhibitory interneurons^15, 16, 19, 20, 22^. Locomotion is also associated with an increase in the gain of visual responses^15, 16, 19, 22^.

Because the most commonly studied active states involve a substantial motor component, it remains unclear whether the associated changes in cortical activity patterns are specific to motor output or more generally attributable to changes in global arousal. Recordings during manipulations of the visual environment suggest that much of the change in firing rates during locomotion is consistent with multimodal processing of visual and motor signals^17, 18^. The integration of locomotor and visual signals in V1 may thus represent elements of predictive coding or play a role in spatial navigation. However, locomotion-associated changes in cortical activity have been replicated by noradrenergic and cholinergic manipulations in the absence of motor output^16, 20, 28^. Changes in V1 activity during locomotion may therefore result from recruitment of neuromodulatory systems that regulate global arousal levels.

Wakefulness comprises states of low and high arousal, but the relationship between changes in arousal and cortical activity remains poorly understood. The functional impact of motor feedback signals to sensory cortex is likewise only beginning to be explored^13, 21, 29, 30^. Here we used behavioral state monitoring and manipulation to dissociate the roles of locomotion and arousal in regulating neural activity in mouse V1. We recorded from V1 in mice head-fixed on a wheel to measure locomotion and monitored pupil diameter to assess arousal^22, 31–33^. To disentangle the effects of motor activity and arousal, we first took advantage of naturally occurring state transitions where mice initiated or finished a bout of locomotion. We examined the precise time course of changes in the LFP, single-unit firing rates, and pupil diameter around locomotion onset and offset. We further compared epochs of high and low arousal during quiescent states. In a second set of experiments, we causally manipulated behavioral state and induced a shift from low to high arousal in the absence of locomotion by delivering an air puff to the animal’s body. We find that arousal mediates most state-dependent changes in LFP activity, whereas increases in overall firing rates are attributable to locomotion. In contrast, enhancement of visual encoding during locomotion is associated with increased arousal, rather than motor activity.

## RESULTS

### Dissociating locomotion and arousal

To separate the contributions of locomotion and arousal to V1 activity and visual encoding, we performed simultaneous recordings of isolated single units and LFPs from multiple sites throughout layers 2-6 of V1 in awake mice (**Fig. 1;** n=88 sessions in 28 mice). Mice were head-fixed on a spring-mounted wheel apparatus (**Fig. S1a**) and recordings were made during both baseline and visual stimulation periods. Behavioral state was assessed by continuous monitoring of arousal and locomotion. To monitor arousal, we measured pupil dilation in the eye that was contralateral to the visual stimulus (**Fig. 1a, Supplementary Movie 1)**. Transition points were defined as shifts from quiescence to locomotion or from locomotion to quiescence and were identified by detecting significant changes in the statistics of the locomotion speed signal from the wheel with high temporal resolution (**Fig. 1b**).

**Figure 1.**
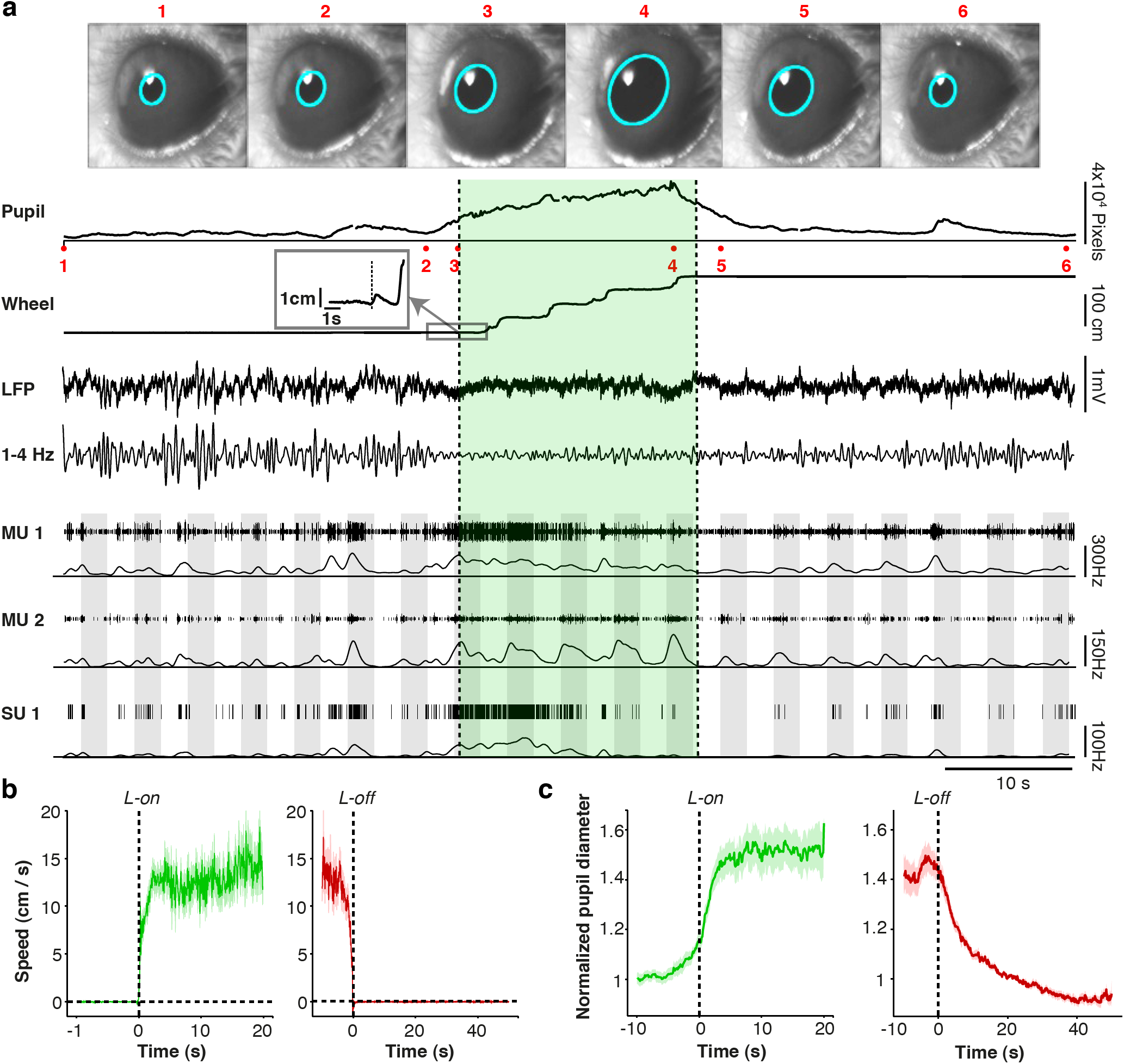
Experimental paradigm to separate the contributions of arousal and locomotion to neural activity and visual encoding in V1. (**a**) Example data from one experiment session. Video frame images of the mouse’s eye (1-6) are shown where acquired at the times indicated in the pupil recording trace. Pupil diameter was recorded on video and extracted posthoc via a fitted ellipse (cyan). The average pupil diameter in pixel units is shown as a function of time. Locomotion is shown as a linearized version of the wheel position. Locomotion onset point is shown in the inset panel. The locomotion period is indicated by green shading. LFP recording is shown as a raw broadband LFP signal for a superficial electrode, together with the 1-4 Hz filtered signals. Thresholded multi-unit traces and spike densities (1 s Gaussian smoothing kernel with SD [Standard Deviation] of 0.25 s) are shown for a superficial (MU1) and deep (MU2) electrode, respectively, together with a single unit trace that was isolated from MU1. Grey shadings indicate visual stimuli at 100% contrast and varying orientations. (**b**) Locomotion speed around locomotion onset (green) and offset (red), shown as mean ± s.e.m (across sessions). (**c**) Population average pupil diameter, normalized to PD at 20-25 s point after locomotion offset, as a function of time around locomotion onset (green) and offset (red), shown as mean ± s.e.m.

We found complex temporal relationships between pupil diameter, locomotion, LFP dynamics, and firing rates of V1 neurons (**Fig. 1a**). At locomotion onset, speed rapidly increased, reaching a plateau after about 2.5 s and remaining at ∼10-15 cm/s until just prior to locomotion offset (**Fig. 1b**). Locomotion speed and pupil diameter were consistently correlated at transition points. Pupil diameter increased prior to the onset of locomotion, suggesting that arousal reliably preceded movement. Pupil diameter reached a plateau after about 2.5 s, and remained elevated until locomotion offset (**Fig. 1c).** Locomotion offset was followed by a gradual decrease in pupil diameter that did not reach baseline values until 40 s later (**Fig. 1c**), indicating a substantial period of elevated arousal in the absence of locomotion. Periods of high arousal and locomotion thus occurred both together and separately, allowing us to discriminate their roles in V1 activity.

### LFP modulation by behavioral state transition

To dissociate LFP changes associated with locomotion and arousal, we computed time-frequency representations of LFP signals around locomotion onset and offset in the absence of visual stimuli. Locomotion onset was preceded by a sharp increase in LFP gamma oscillations (55-65 Hz) and a decrease in low-frequency LFP fluctuations (1-4 Hz; **Fig. 2a**). Gamma power decreased gradually over time after locomotion onset, whereas low-frequency fluctuations remained suppressed throughout the locomotion period (**Fig. 2a**). In contrast, locomotion offset was followed by a gradual increase in low-frequency LFP power and a gradual decrease in LFP gamma power that lasted up to 40-50 s (**Fig. 2b**). The time courses of LFP low-frequency and gamma power were strongly correlated (Spearman correlation) with the time course of pupil dynamics, but not with the time course of locomotion speed (**Fig. 2c**). The correlations between LFP power and pupil diameter time courses were strongly linear (gamma and pupil: Pearson’s R = 0.94, p=2.4×10^−6^; 1-4 Hz and pupil: Pearson’s R = −0.93, p=4.1×10^−5^), suggesting that arousal, rather than locomotion, mediates most of the observed change in LFP patterns.

**Figure 2.**
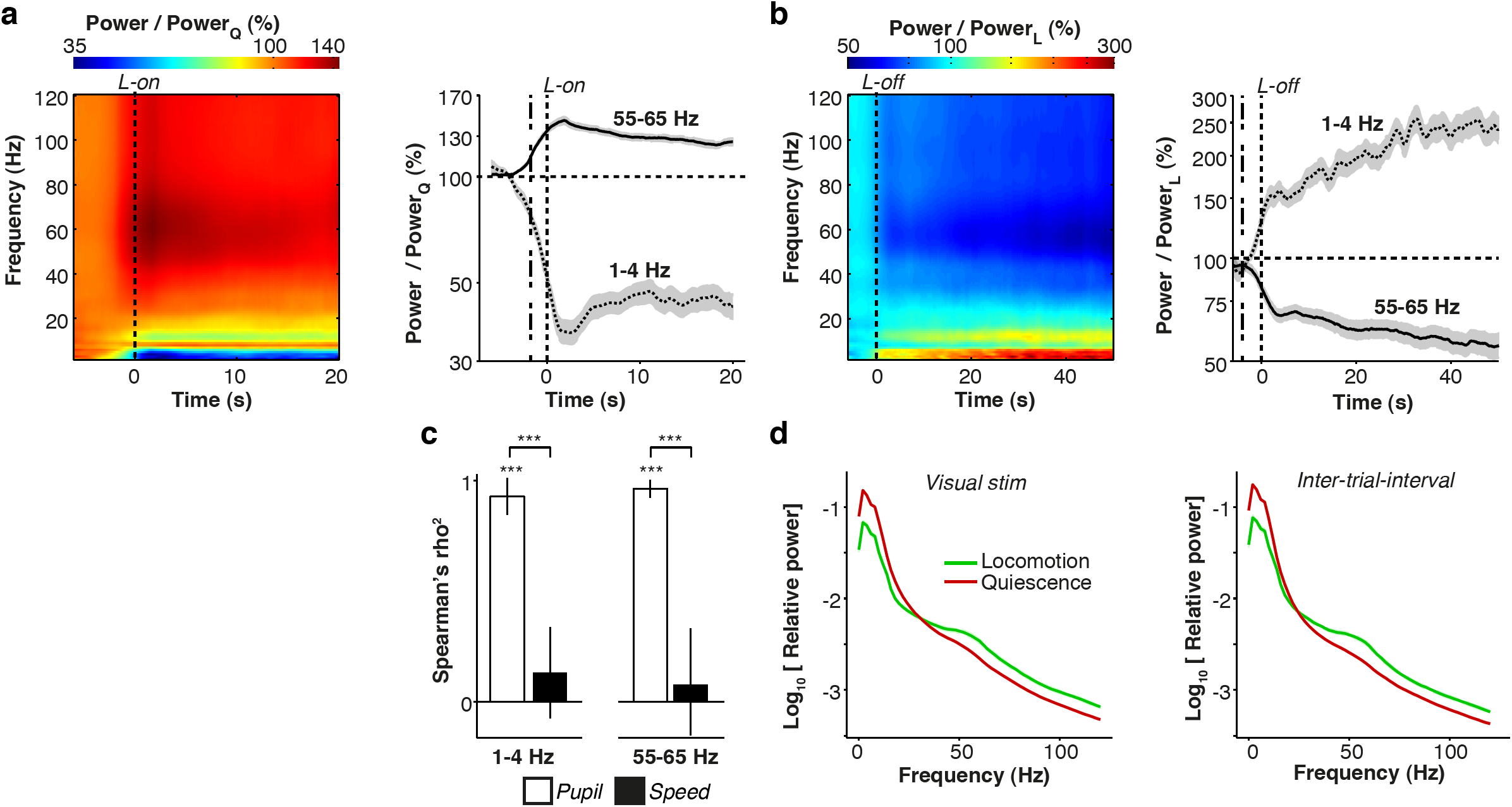
Contributions of arousal and locomotion to V1 LFP rhythms. (**a**) Left: Time-frequency representation of LFP signals around locomotion onset (L-on), showing base-10 log-transformed relative power, i.e. power at time *t* divided by average power in the [−4.5,−4] s quiescent interval preceding L-on. Power was computed in sliding windows of ±2 s using short-term Fourier Transform. Log power ratios are shown as percentages. Right: Line plots of gamma (55-65 Hz) and low-frequency (1-4 Hz) power across n=58 sites in 11 mice, shown as mean ± s.e.m. Dashed lines at 2 s before zero represent the effective time at which data after L-on would be included in computation, revealing spectral changes before this point. Note the appearance of an 8-10 Hz theta-band that interrupts the reduction in low-frequency LFP fluctuations, presumably due to volume conduction from the hippocampus. (**b)** Time-frequency representation of V1 LFP signals around locomotion offset (L-off). Relative power was computed by dividing by the average power in the [−3,-2.5] s epoch before locomotion offset, for n=133 sites in 18 mice. (**c**) Spearman’s correlation (rho^2^) between either average pupil diameter (PD) (open bars) or locomotion speed (closed bars) with average gamma and delta time courses for quiescence period, using data >−3 s before locomotion offset till 40 s after locomotion offset.***: p<0.001. Shown mean ± s.e.m. n=133 sites in 18 mice. (**d**) Average power spectra during visual stimulation period (left) and ITI (right), separately for locomotion and quiescent periods, for n = 196 sites in 23 mice. Power spectra were computed using multitapering of 500 ms windows, with a smoothing window of ±4 Hz.

We also recorded LFP activity in V1 while presenting visual stimuli. All stimuli were drifting gratings on a mean luminance background (**see Online Methods**). We found that the increase in LFP gamma-band power and decrease in low-frequency power during locomotion periods were observed for both visual stimulation and inter-trial interval (ITI) epochs (**Fig. 2d**). Increased gamma oscillations during locomotion are thus largely the result of behavioral state transition, rather than visual stimulation.

### Cell-type specific modulation of firing rates

Next, we investigated the contributions of locomotion and arousal to V1 firing rates in the absence of visual stimuli. We compared firing rates between epochs of locomotion and quiescence and within each type of epoch to examine the temporal dynamics of changes in firing around transition points (**Fig. 3a**). Individual V1 neurons demonstrated a broad range of relationships between spontaneous firing rate and behavioral state (**Fig 3a, Fig. S2**). To characterize firing patterns across the population, we classified well-isolated single units into 34 fast spiking (FS), putative inhibitory, and 157 regular spiking (RS), putative excitatory cells based on action potential waveform characteristics (**Fig**. **S3**).

**Figure 3.**
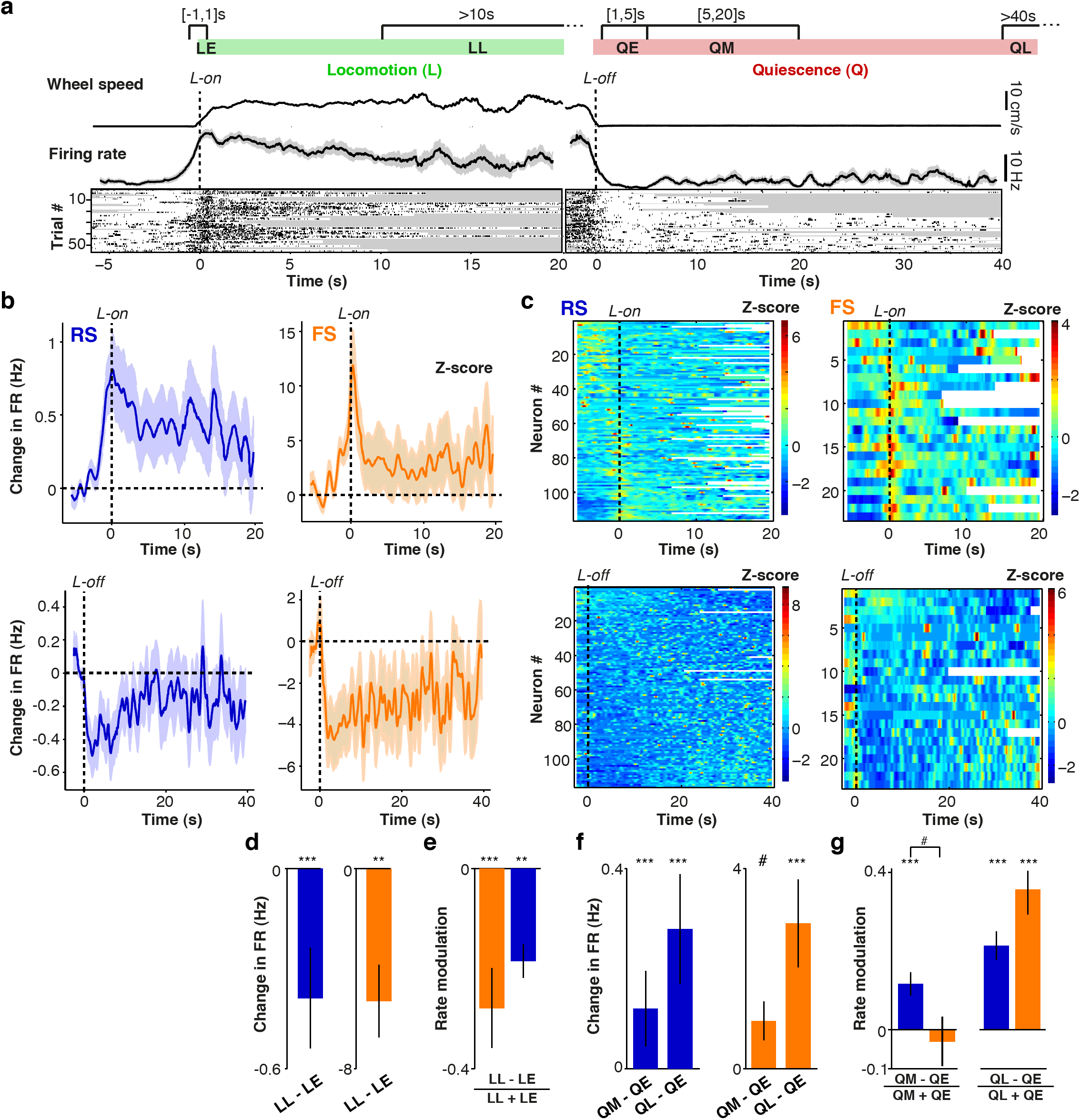
Contributions of arousal and locomotion to spontaneous firing activity in V1. **(a)** Schematic showing division of data into epochs for analysis. *LE*: early locomotion period (-1 to 1 s around L-on). *QE*: early quiescent period (1 to 5 s after L-off). *QM*: middle quiescent period (5 to 20 s after L-off). *QL*: late quiescent period (>40 s after L-off). Plots show average wheel speed and firing rate for an example FS cell during locomotion onset (L-on, left) and locomotion offset (L-off, right). Rasters show individual transition points for the same cell. Gray shadings indicate L-off and L-on during individual trials. (**b**) Upper: Population average change in RS (blue) and FS (orange) cells’ firing rates relative to a quiescent baseline period (taken −6 to −3 s before L-on) as a function of time around L-on, shown as mean ± s.e.m. Lower: Change in firing rates relative to a baseline locomotion period (defined −3 to −1 seconds before L-off) as a function of time around L-off. (**c**) Upper: Overview of all recorded RS (left) and FS (right) cells, sorted by firing rate difference between quiescent baseline and the [-1,1] s interval around L-on. Color coding corresponds to Z-scored firing rates. Lower: As in the upper panel, but as a function of time around L-off and sorted on Spearman’s correlation between time after L-off (>3 s) and quiescence. (**d-e**) Mean ± s.e.m. of firing rate differences (**d**) and firing rate modulation (**e**) between LE and LL epochs. Statistical testing using two-sided Rank-Wilcoxon tests. #,*,**,*** correspond to p<0.1, 0.05, 0.01, 0.001. RS: n=106/17, FS: n=18/9. (**f-g**) As in (**d-e**), but now for QE vs. QL and QM vs. QL. RS: 101/18, 101/18 (#cells/#mice), FS: 21/10, 18/9. Statistical comparison of QM vs. QL rate modulation: RS, FS: p=7.1×10^−4^, 8.6×10^−4^.

Both FS and RS cells’ firing rates increased before locomotion onset, peaked around the time of locomotion onset, and declined over time throughout the locomotion period (**Fig. 3b-c**). To quantify the anticipatory effect for individual cells, we computed the Pearson cross-correlation coefficient between locomotion speed and instantaneous firing rate in the interval around locomotion onset. Most cells exhibited a strong linear correlation between locomotion speed and firing rate that was forward-shifted in time, indicating that increases in RS and FS firing rates preceded increases in locomotion speed (**Fig. S4**).

Both FS and RS firing rates were significantly higher during the early period around locomotion onset (within ±1 s of locomotion onset; LE) than during the late locomotion period (>10 s after locomotion onset; LL) (**Fig. 3d, Fig. S5a**). To directly compare state-dependent changes between RS and FS cells, we computed a rate modulation value (FR_b_ – FR_a_**) /** (FR_a_ + FR_b_), which normalizes for absolute rate differences. RS and FS cells exhibited similar degrees of rate modulation during the early vs. the late locomotion period (p=0.48; **Fig. 3e**).

We next examined whether increases in firing rate during locomotion might be explained by the associated change in arousal. To separate the effects of locomotion and arousal, we examined the trajectories of firing rates during quiescence following locomotion offset, when pupil diameter showed substantial variation in the absence of movement (**Fig. 1b-c**). For both RS and FS cells, locomotion offset was accompanied by a rapid decrease in firing rates, followed by a subsequent gradual increase over time (**Fig. 3b**).

To directly compare firing rates across epochs within the quiescent period, we analyzed early quiescence (1-5 s after locomotion offset; QE), middle quiescence (5-20 s after locomotion offset; QM) and late quiescence (>40 s after locomotion offset; QL) (**Fig. 3a**). Pupil diameter was significantly decreased across the QE to QM to QL intervals after locomotion offset (**Fig. S6**). In contrast, FS and RS firing rates significantly increased across the QE to QM to QL intervals after locomotion offset **(Fig. 3f-g)**. We found no significant difference in rate modulation between FS and RS cells across quiescent epochs **(Fig 3f-g**, **Fig. S5a**).

To further quantify the trajectory of V1 firing rates over time, we computed Spearman correlation coefficients between firing rate and time after locomotion offset (using only data >3 s after locomotion offset; **Online Methods**). Many cells showed a significant association between firing rate and time (FS: 10/23, RS: 37/80, at p<0.05), and the average correlation was higher than zero (**Fig**. **S5b**, mean ± s.e.m. of Spearman’s rho = 0.04±0.024, p=0.034; Rank-Wilcoxon test). However, we also found subpopulations of cells whose firing rate either decreased or increased over time (**Fig. 3c**, **Fig. S5b**, FS positive: 6/23, negative: 4/23; RS positive: 20/80, negative: 17/80, p<0.05), suggesting diverse firing rate trajectories across the V1 population.

Together, these results suggest that both RS and FS cells in V1 demonstrate increased spontaneous activity in anticipation of and during locomotion. However, a state of heightened arousal, as indicated by maintained pupil dilation after the cessation of movement, is accompanied by decreased RS and FS firing. In contrast to the spectral changes in LFP signals, elevated firing during movement is thus specific to locomotion periods and unlikely to result from associated changes in global arousal.

### Modulation of visual encoding by state transition

To determine the impact of locomotion and arousal on visual encoding, we recorded LFP and unit activity in V1 while presenting visual stimuli. Visual stimulation during locomotion caused an increase in gamma band power at frequencies above 60 Hz, with a spectral peak around 75 Hz (**Fig. S7a-b**). During quiescent periods, visual stimulation instead caused an increase in gamma-band power at a broader range above 20 Hz, with a relatively shallow spectral peak around 30 Hz (**Fig. S7a-b**). We further found that contrast tuning of LFP gamma-band power was affected by locomotion, with increased and decreased gain at high and low gamma frequencies, respectively (**Fig. S7c-f**).

The robust differences in visually evoked LFP dynamics between periods of locomotion and quiescence suggest that visual stimulation may likewise differentially affect firing rates during these states. We therefore computed the mean peak firing rate of RS cells in response to stimuli presented during locomotion and quiescence **(Fig 4a-b)**. To separate the effects of locomotion and arousal, we divided the quiescence period into separate epochs as in **Figure 3**. The exact durations of the epochs were adjusted to account for the duration and timing of stimulus presentation. We analyzed early quiescence (3-20 s after locomotion offset; QE), when the pupil was relatively dilated, and late quiescence (>40 s after locomotion offset; QL), when the pupil was relatively constricted. We used the signal-to-noise (SNR) ratio, defined as (FR_stim_ - F_RITI_) / (FR_stim_ + FR_ITI_), to assess visual encoding by V1 neurons during each epoch (**Online Methods**). We found an overall increase in the SNR of visual encoding during locomotion in comparison to quiescence for RS, but not FS, cells (**Fig. 4c**). A finer analysis of time periods found that the SNR in early quiescence (QE), during high arousal, was larger than in late quiescence (QL), during low arousal, for both RS and FS cells (**Fig. 4c**). We found that spontaneous, but not visually evoked, RS firing rates were significantly higher in the late than in the early quiescence period (**Fig. S8a-b**). The observed increase in SNR during high arousal states thus largely resulted from the differential effects of arousal on spontaneous and evoked firing rates.

**Figure 4.**
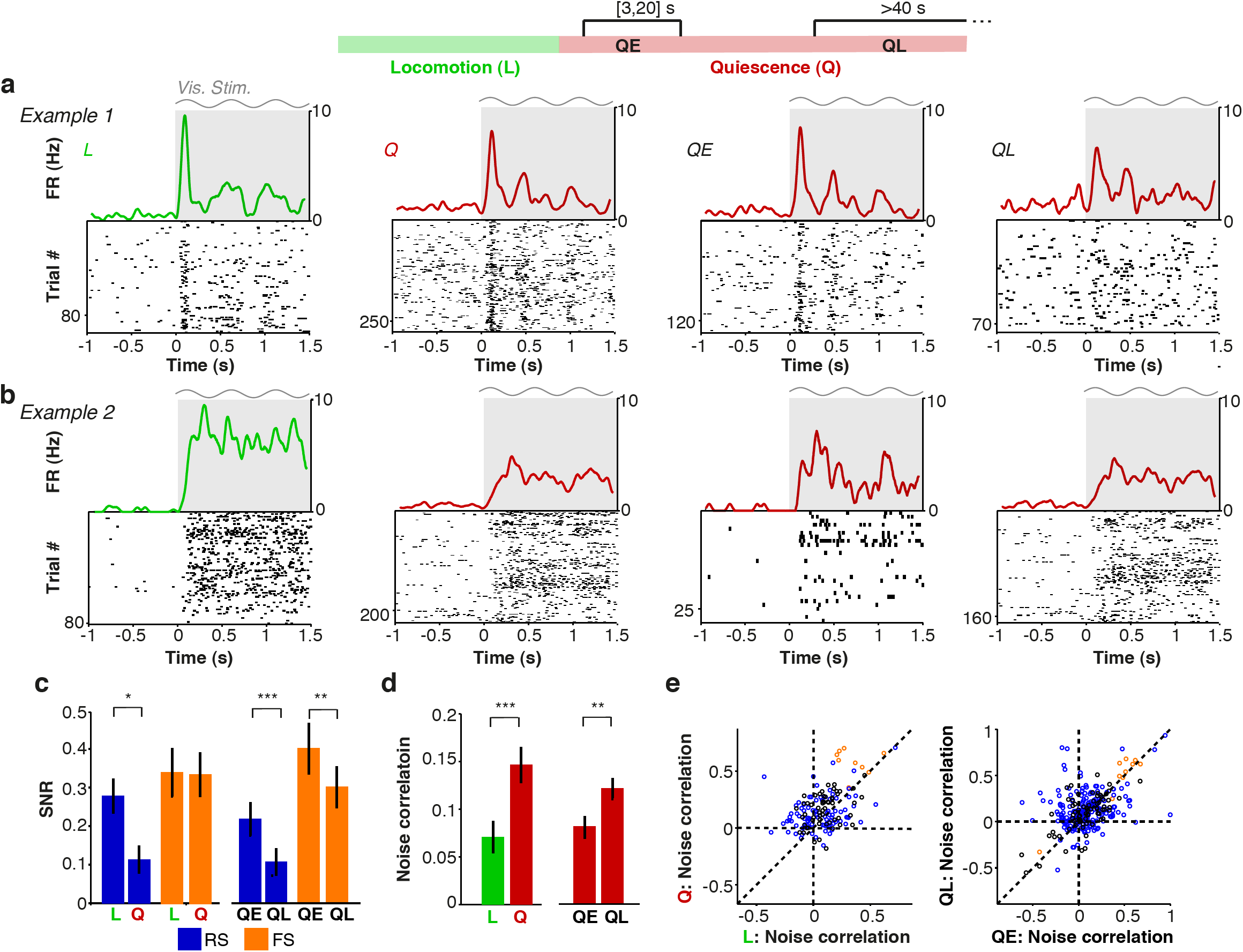
Contributions of locomotion and arousal to visual encoding and noise correlations in V1. (**a,b**) Raster plots of the visual responses of two example RS cells with associated firing rate density (computed using ±0.025 s Gaussian kernels with SD of 0.0125 s) for all locomotion (L), all quiescence (Q), early quiescence (QE; 3 to 20 s after locomotion offset) and late quiescence (QL; >40 s after locomotion offset). Gray shading and sinusoid indicate visual stimulation. (**c**) Mean ± s.e.m. of signal-to-noise ratio (SNR), defined as (FRstim-FRITI) / (FRstim+FRITI), for each period. #,*,**,*** correspond to p<0.1, 0.05, 0.01, 0.001, two-sided Rank-Wilcoxon test. RS: n=88/20, 101/20 (#cells/#mice). FS: n=23/10, 24/11. (**d**) Mean ± s.e.m of noise correlation of firing rates during different behavioral periods. n=155/9, 289/11 (#pairs/#mice). *,**,***: p<0.05, p<0.01, p<0.001, two-sided Rank-Wilcoxon test. (**e**) Left: Noise correlation for locomotion vs quiescence. Circles correspond to cell pairs. FS-FS: n=9/3, RS-RS: 90/7, FS-RS (black): n=56/5. Right: as left, but for QE vs. QL. FS-FS: 10/4, RS-RS: 206/7, FS-RS: n=73/7.

The SNR is a measure of the encoding of visual stimuli by individual neurons. However, visual stimuli are likely to be encoded by patterns of V1 population activity^34, 35^. We therefore calculated noise correlations, which may determine the efficiency of population coding^36, 37^, for each epoch (**Online Methods**). Overall, noise correlations were significantly lower during periods of locomotion than during quiescence (**Fig. 4d-e**). Because low-frequency and gamma-band LFP power showed gradual changes over time after locomotion offset, we hypothesized that noise correlations would increase with time after locomotion offset. Indeed, noise correlations were significantly elevated during late quiescent periods, when arousal was low, as compared to early quiescent periods (**Fig. 4d-e**).

These results suggest that visual encoding is enhanced during locomotion as a result of increased SNR and decorrelation of visually evoked activity. However, these changes are mainly attributable to arousal, rather than locomotion, as comparable changes in SNR and noise correlations were observed during periods of heightened arousal in the absence of movement.

### Isolating the cortical effects of arousal

The analyses of naturally occurring state transitions described above suggest separable effects of locomotion and arousal on firing rates, LFP activity, and visual encoding in V1. However, it is possible that the period of heightened arousal following locomotion offset represents an unusual form of arousal or is affected by long-lasting modulation of V1 activity by motor signals. To more causally test the role of arousal in regulating neural activity, we induced a state of enhanced arousal by puffing air on the back of the mouse (**Fig. S1b**, **Online Methods**). In a subset of sessions (n=20 sessions in 10 mice), we administered brief air puffs to the mouse while monitoring pupil diameter and performing simultaneous recordings of V1 LFPs and isolated single cells (**Fig. 5**; n = 61 cells in 10 mice).

**Figure 5.**
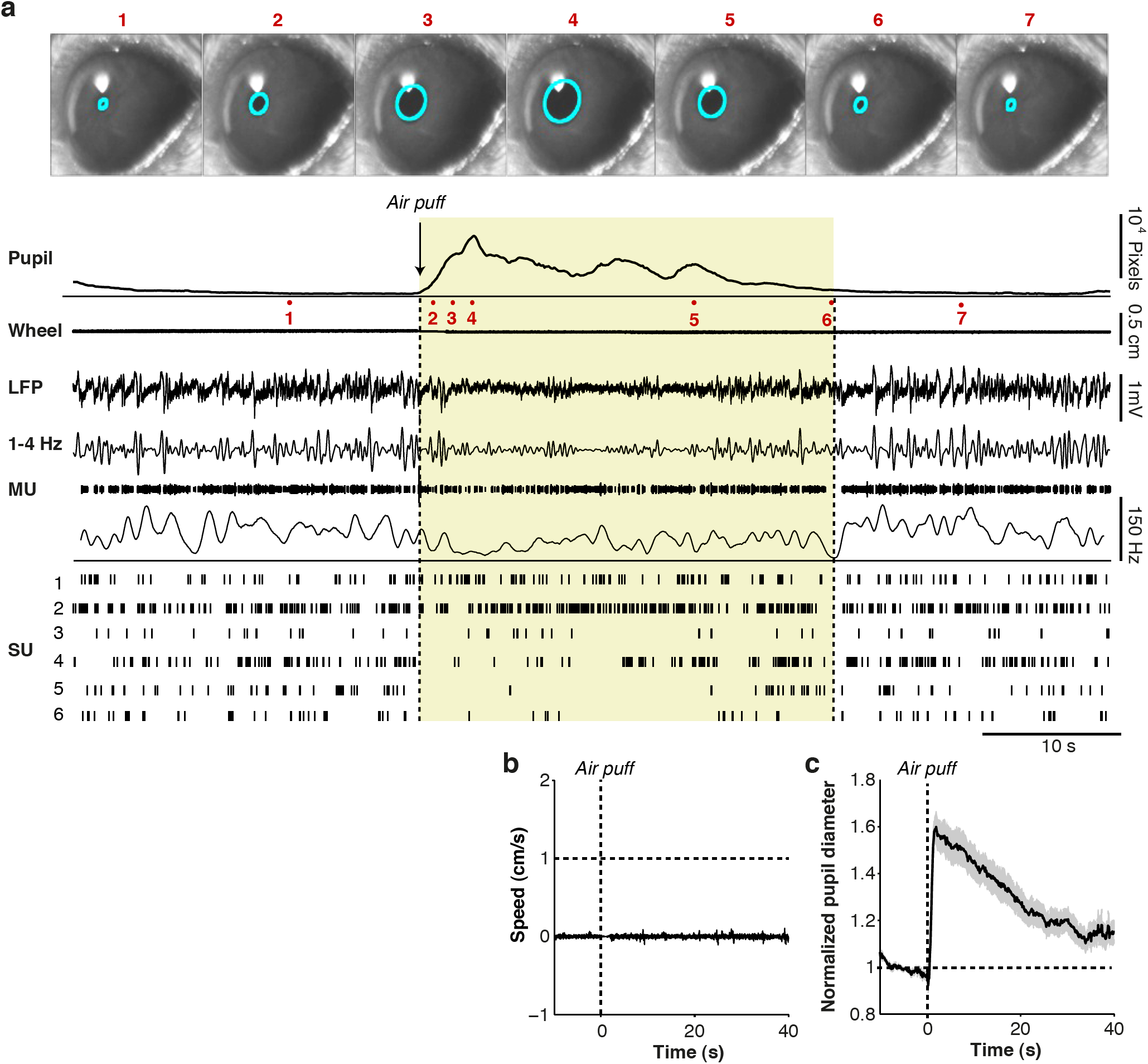
Causal induction of arousal without locomotion. (**a)** Example data from one experiment session. Video images captured of eye are shown for several time points (1-7). Average pupil diameter trace in pixels is shown as a function of time around air puff onset. Locomotion is shown as a linearized version of the wheel position. LFP recording is shown for one electrode, together with the 1-4 Hz filtered signal. A multiunit trace (MU) and associated spike density plot are shown for the same electrode. Single-unit traces (SU1-6) were isolated from a total of three electrodes in L5/6 during this recording session. Shaded yellow area indicates period during which pupil was dilated, for visualization purposes. (**b**) Average locomotion speed as a function of time around the air puff. (**c**) Average pupil diameter, normalized to pre-air-puff interval (-10 to 1 s before air puff), as a function of time. The s.e.m.’s were computed across sessions (n=20 in 10 mice).

Delivery of the air puff reliably induced both arousal, as measured by pupil dilation, and changes in unit activity in the absence of locomotion (**Fig. 5a, Supplementary Movie 2**). Across sessions (n=20 sessions in 10 mice), air puffs caused an average 1.6-fold increase in pupil diameter in the absence of locomotion (**Fig. 5b-c**). This increase was similar in magnitude to the average increase in pupil diameter observed during locomotion (**Fig. 1c**), suggesting a comparable level of arousal.

### LFP modulation by induced arousal

Our analysis of arousal periods following locomotion offset (**Fig. 2**) predicts that a shift to an aroused state in the absence of locomotion should be accompanied by a decrease in low-frequency LFP power and an increase in gamma-band power. Indeed, we found that after the air puff the LFP showed a long-lasting, 2.5 fold decrease in low-frequency fluctuations comparable in magnitude to the locomotion effect (**Fig. 6a-b**). The induced arousal also caused a 1.25-fold increase in gamma-band power. The pupil diameter time course showed a strong positive correlation with gamma-band power, and a negative correlation with low-frequency power (**Fig. 6c**). These findings indicate that a shift to a state of heightened arousal without locomotion causes a decrease in low-frequency fluctuations and an increase in gamma-band fluctuations in V1.

**Figure 6.**
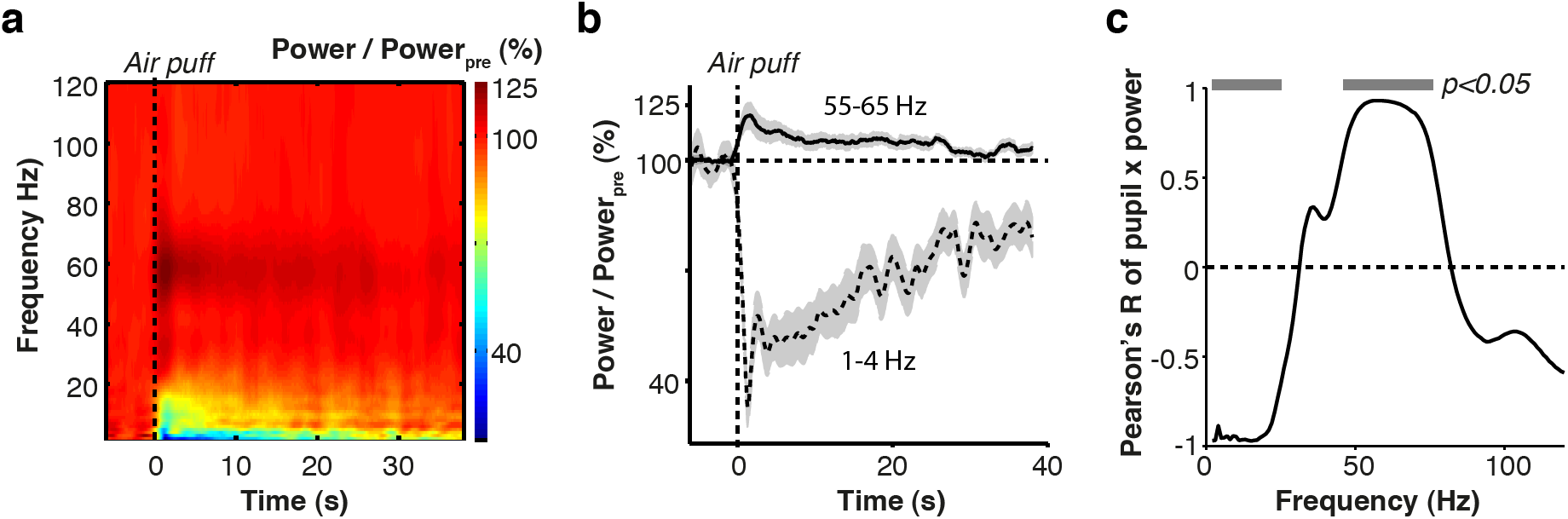
Arousal causes a frequency shift in V1 LFP activity. (**a**) Time-frequency representations of LFP signals around the air puff. Base-10 log-transformed relative power, i.e. power at time *t* divided by average power in the [-6,-1] s interval preceding the air puff. n = 45 sites/10 mice. (**b**) Line plots of gamma and delta power, with shading representing s.e.m. across sites, around time of air puff. **(c)** Correlation of pupil diameter time course with LFP time course at various frequencies. Gray horizontal bars indicate significance at p<0.05, linear regression analysis.

### Rate modulation by induced arousal

We next investigated how firing rates were affected by the causal manipulation of arousal levels. Overall, we found that the air puff caused a significant, long-lasting decrease in spontaneous firing rates **(Fig. 7a,b)**. However, a small subpopulation of RS and FS cells showed elevated firing rates after the air puff (**Fig. 7a,c**). Air puffs were particularly suppressive for RS cells if they had a high propensity to engage in irregular, burst firing (**Fig. 7d; Online Methods**), but a similar relationship was not observed for FS cells (**Fig. 7d**).

**Figure 7.**
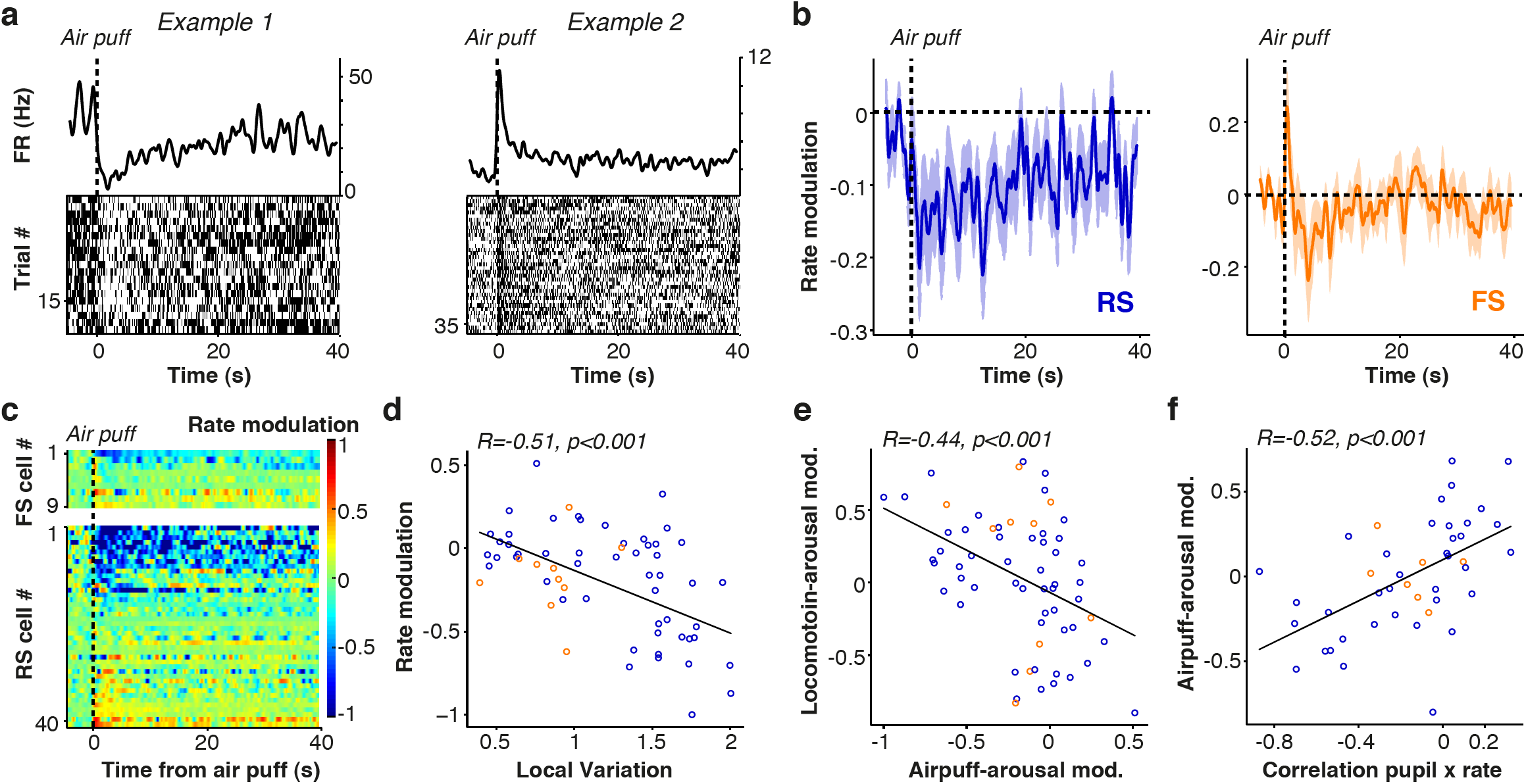
Arousal suppresses firing rates in V1. (**a**) Raster plots and spike density traces (1 s windows with Gaussian kernel and SD of 250 ms) from two example (left: FS, right: RS) cells are shown as a function of time. (**b**) Average modulation of firing rate induced by the air puff for RS (blue) and FS (orange) cells. Mean ± s.e.m of modulation in [1,4] s interval for RS = −0.19±0.046, p=4.6×10^−4^, n=51/9 (#cells/#mice). FS: −0.14±0.074, p=0.084, n=10/6. Difference FS and RS cells not significant (Rank-Wilcoxon test; p=0.87). (**c**) Rate modulation scores for all FS (upper) and RS (lower) cells as a function of time, relative to 6 s prior to air puff. (**d**) Air puff modulation index, comparing [1,4] s post air puff to 6 s prior to air puff, as a function of local variation coefficient, a measure of firing regularity. High local variation values correspond to bursty cells and a local variation coefficient of 1 indicates Poisson-like firing (**Online Methods**). RS cells: R = −0.57, p=1×10^−5^, n=51/9 (#cells/#mice), FS cells: R=0.13, p=0.71, n=10/6. (**e**) Values on the x-axis indicate air puff modulation index. Values on the y-axis indicate airpuff-arousal modulation, calculated as the Spearman correlation of time after locomotion (>3 s) and firing rate. R = −0.47, p = 4.15×10^−4^, NS: R = −0.32, p=0.36, difference RS and FS n.s., Fisher Z-test. (**f**) Air puff modulation index vs. correlation of pupil diameter and firing rate across trials. Correlation of pupil and firing rate was defined over trials using the [1,4] s interval after the air puff. Pearson’s R = 0.52, p=3.3×10^−4^, n=45/7.

Individual neurons should show similar effects in response to naturally occurring and induced arousal. We therefore predicted that cells whose firing rates were suppressed by the induced arousal would likewise show suppression after locomotion offset, followed by gradual increases in firing rate over time as arousal levels decreased (**Fig. 3**). For this analysis, we selected only locomotion epochs that did not occur within 50 s of an air puff. We first defined a *locomotion-arousal modulation* as the correlation between time after locomotion offset and firing, with positive values indicating that a cell was suppressed by arousal. We also defined the *airpuff-arousal modulation* in the interval after the air puff. We found that the *locomotion-arousal* and *airpuff-arousal* modulations were significantly negatively correlated, demonstrating consistent effects of arousal on firing rate (**Fig. 7e**).

The coordinated effects of induced arousal on firing rate and pupil diameter also predict that fluctuations in arousal should correlate on a trial-by-trial basis with firing rates. We predicted (more) negative correlations between pupil diameter and firing rates across trials for cells whose firing was on average suppressed by the air puff. Likewise, we predicted (more) positive correlations for cells whose firing was enhanced by the air puff. These predictions were supported by trial-by-trial correlation analysis (**Fig. 7f**). Together, these results suggest that global arousal is associated with decreased spontaneous firing rates.

### Arousal and visual encoding

To isolate the role of arousal in modulating single cell and population visual encoding, we administered air puffs randomly in combination with presentation of visual stimuli in a subset of experiments (**Online Methods**). We then compared visual responses in the 10 s before the air puff with those in the 10 s after the air puff. Across the population of cells, we found a significant increase in SNR following the induced arousal (**Fig. 8a,b**). We also observed that induced arousal significantly decreased noise correlations (**Fig. 8a,c**). These results indicate that arousal alone replicates the effects of locomotion on visual encoding, leading to an increase in the salience of visual responses by individual cells and decorrelating activity across the population.

**Figure 8:**
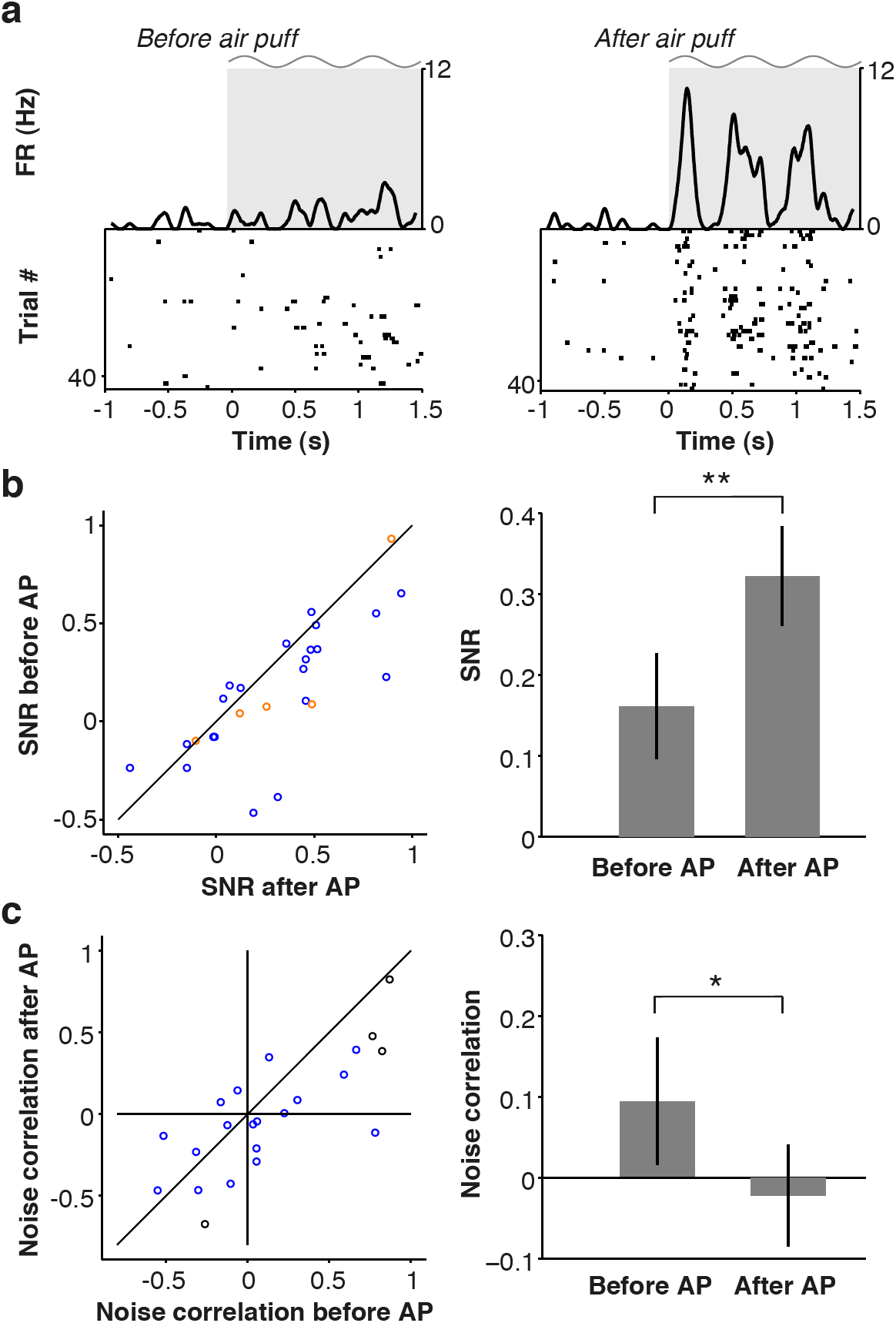
Arousal enhances visual encoding and decreases noise correlations in V1. (**a**) Raster plots of the visual responses of an example cell with associated firing rate density (computed using ±0.025 s Gaussian kernels with SD of 0.0125) before (left) and after (right) air puff (AP). Gray shading and sinusoid indicate visual stimulation. (**b**) Plot of signal-to-noise ratio (SNR) for 10 s interval before air puff (y-axis) compared to 10 s interval after air puff (x-axis). Population SNR shown at right, as mean ± s.e.m (n=25/8, #cells/#pairs). (**c**) Noise correlation in the 10 s after air puff compared to the 10 s before air puff. Circles correspond to cell pairs (red = FS, blue = RS, black = FS/RS). Population average noise correlation is shown at right. (**b-c**) *, **: p<0.05, p<0.01, n=22/4 (#pairs/#mice), two-sided Rank-Wilcoxon test.

## Discussion

We examined the distinct contributions of arousal and locomotion to V1 activity and visual encoding using extracellular recording in combination with behavioral state monitoring and manipulation. Our data revealed that much of the change in V1 LFP and unit activity associated with locomotion is the result of increased arousal. We found that heightened arousal in the absence of locomotion has several cortical impacts: 1) a change in the temporal patterning of neural activity, comprising a reduction in low-frequency oscillations and an increase in gamma synchronization, 2) a net suppression of firing rates, and 3) a change in visual encoding at both the single-cell and population levels. In contrast, locomotion specifically contributes to an increase in RS and FS cell firing rates in anticipation of and during movement. Arousal state and motor activity thus have distinct roles in regulating activity in primary visual cortical circuits.

We found substantial differences in V1 activity between periods of quiescence and locomotion. This is consistent with several studies that used electrophysiology or calcium imaging in awake head-fixed rodents^15–20, 22, 28^. However, previous results suggested that elevated RS cell firing during locomotion periods was not associated with locomotion, but was rather a modulation of visual responses^15, 19^. By focusing on state transition points, we were able to show that RS firing was strongly increased in anticipation of locomotion and during movement, even in the absence of visual stimulation. In contrast, causal induction of a state of high arousal caused a suppression of firing rates, indicating that motor and arousal signals may have opposing effects on spontaneous spiking activity in V1. Studies in auditory and barrel cortex have found that firing rates of both RS and FS cells are suppressed by locomotion and whisking, respectively^12, 14, 21, 38^. Thus, increased RS and FS firing during locomotion is likely specific to visual cortex and may be related to the necessity for integration of motor and visual signals in order to faithfully represent the outside world with respect to the animal’s position^17, 18^.

Firing rates and LFP power in the low (1-4Hz) and gamma (55-65Hz) ranges showed changes well in advance of the onset of locomotion, as did pupil diameter. These anticipatory changes indicate that increased arousal levels and motor-related activity in V1 reliably precede the execution of motor output. Increased firing in anticipation of locomotion, consistent with motor planning or predictive coding^17, 18^, could rely on top-down inputs from fronto-parietal circuits, in which many cells fire in anticipation of saccades and other movements^39, 40^. Indeed, recent work has highlighted the existence of long-range inputs from frontal to striate cortex^41^ and motor cortical areas may also project to V1 or other visual areas^42, 43^.

Distinct V1 cell populations exhibited different modulation around transition points. We observed enhanced signal-to-noise during locomotion for visual responses only in RS, putative excitatory cells but not in FS, putative inhibitory cells, potentially indicating cell-type specific regulation of visual encoding. Recent work has highlighted distinct patterns of state-dependent recruitment of different interneuron classes^14, 16, 20, 22, 38^. Notably, we found diversity within both the RS and FS cell populations in the trajectory of state-dependent changes in firing patterns at behavioral state transition points. Whereas most RS and FS cells were suppressed by arousal, a subset of both cell types instead showed enhanced firing. Firing rate suppression with arousal was particularly evident in bursting RS cells.

We took advantage of direct manipulations of behavioral state to probe the dynamic range of behavioral state-dependent cortical activity. Causal induction of arousal with the air puff stimulus initiated a shift from low to high frequencies in the cortical LFP. We found that induced arousal replicated the 2.5 fold reduction in 1-4 Hz fluctuations observed during locomotion periods. Low frequency fluctuations increased only gradually after locomotion offset, with a strong linear relationship to the pupil diameter. Arousal alone therefore appears to account for the reduction in 1-4 Hz LFP power during locomotion periods. The observed linear relationship between pupil diameter and 1-4 Hz LFP power differs from recent work in which pupil diameter did not correlate with low-frequency membrane potential power, likely because only very small and short-lasting pupil fluctuations were considered^22^. LFP and firing rate changes were dissociated from one another during arousal periods, as the gradual increase in LFP low-frequency power and decrease in LFP gamma-band power were accompanied by a net increase in firing rates over the same time period.

The effect of arousal on gamma-band LFP power in the absence of locomotion (∼20% increase) was smaller than the increase in gamma-band LFP power during locomotion periods (∼40% increase). Gamma-band power showed a similar adaptation during the locomotion period as firing rates, in contrast to 1-4 Hz power. A change in arousal might therefore not exclusively account for the increase in gamma-band power during locomotion periods, suggesting that locomotion might further amplify gamma-band oscillations through the increased drive of local RS and FS cells^44–47^. Previous work has linked noise correlations to intrinsic low-frequency fluctuations and a decrease in gamma-band oscillations^25, 27, 48, 49^. The effects of arousal observed here thus overlap with those of focused spatial attention in primates, where attention is associated with increased temporal patterning in the gamma band and decreased noise correlations^23, 25–27^.

State-dependent changes in visual encoding were highly correlated with arousal level across a wide range, suggesting extensive flexibility in the sensory processing operations of cortical circuits. An increase in the SNR of visual responses in V1 has previously been reported for waking vs. sleeping states^2, 7^, indicating possible regulation of visual sensitivity by overall arousal levels. In agreement with this idea, we found that states of high arousal were associated with increased SNR and decreased noise correlations, indicating enhanced encoding of visual stimuli at both the single-cell and population levels. We found a strong trial-by-trial correlation between arousal, as measured by pupil diameter, and both measures of visual encoding, suggesting a dynamic system in which changes in arousal fine-tune the gain of visual responses in V1 on a moment-to-moment basis.

Lesion and stimulation studies have supported causal roles for several major neuromodulatory systems in controlling sleep-wake transitions and the temporal pattern of cortical activity, including norepinephrine (NE) and acetylcholine (ACh)^28, 50–55^. Like arousal, neuromodulatory action desynchronizes the cortex, promotes gamma oscillations, and changes sensory encoding^4, 20, 25, 51, 53, 54^. Noradrenergic blockade of awake mouse V1 eliminates the depolarization and elevated firing associated with locomotion^16^. Recent evidence also points to the involvement of the mesencephalic locomotor region, a cholinergic brainstem nucleus, in the control of locomotion and regulation of V1 activity patterns and visual responses^28^. In addition, cholinergic afferents from the basal forebrain nucleus of the diagonal band of Broca selectively target V1 interneurons involved in locomotion-related changes in firing rates^20^. Arousal effects observed in V1 during waking state transitions are thus likely involve complex interactions between multiple neuromodulatory systems.

In summary, our data show that activity in mouse visual cortex during wakefulness is differentially regulated by arousal and motor signals. We find a complex interaction between internally generated cortical states and visual inputs. Arousal restructures spontaneous cortical activity and promotes fast gamma-band oscillations, which may be optimal for synchronized, bottom-up routing of sensory signals^23, 48, 56, 57^ This shift in the mode of cortical operation also strongly increases the signal-to-noise ratio of visual representations. The interplay between arousal level, motor activity, and sensory input may contribute to the functional flexibility of cortical circuits.

## Acknowledgements

This research was supported by a NARSAD Young Investigator award, an Alfred P. Sloan Fellowship award, a Whitehall grant, a Klingenstein fellowship award, a McKnight Scholar award, and NIH/NEI grants R00 EY018407 and R01 EY022951 to JAC, a Rubicon Grant (Netherlands Organization for Science) to MV, a Jane Coffin Childs Fund fellowship award to RBB, and a NARSAD Young Investigator award to UK. We thank James Mossner and Hyun Lee for technical support. We also thank the main developers of M-Clust (D. Redish), KlustaKwik (K. Harris) and FieldTrip (R. Oostenveld) for free use of their software. We thank M.J. Higley for comments on the manuscript.

### Author contributions

MV, RBB, UK and JAC designed research. RBB, MV and JAC conducted experiments. UK and JAC developed hardware. MV analyzed data. MV and JAC wrote the manuscript. MV, RBB, and UK contributed equally to this study.

## Online Methods

### Animals

All animal handling was performed in accordance with guidelines approved by the Yale Institutional Animal Care and Use Committee and federal guide. All mice were 4-6 months old wild type males and housed 3-5 per cage on a 12h light-dark cycle.

### Headpost surgery and wheel training

Mice were handled for 5-10 min/day for 5 days prior to the headpost surgery. On the day of the surgery, the mouse was anesthetized with isoflurane and the scalp was shaved and cleaned three times with Betadine solution. An incision was made at the midline and the scalp resected to each side to leave an open area of skull. Two skull screws (McMaster-Carr) were placed at the anterior and posterior poles. Two nuts (McMaster-Carr) were glued in place over the bregma point with cyanoacrylate and secured with C&B-Metabond (Butler Schein). The Metabond was extended along the sides and back of the skull to cover each screw, leaving a bilateral window of skull uncovered over primary visual cortex. The skin was then glued to the edge of the Metabond with cyanoacrylate. Analgesics were given immediately after the surgery and on the two following days to aid recovery. Mice were given a course of antibiotics (Sulfatrim, Butler Schein) to prevent infection and were allowed to recover for 3-5 days following implant surgery before beginning wheel training.

Once recovered from the surgery, mice were trained with a headpost on the wheel apparatus. The mouse wheel apparatus was 3D printed (Shapeways Inc.) in white plastic with a 15 cm diameter and integrated axle and was spring-mounted on a fixed base. The initial wheel design was adapted from ^58^ (**Fig. S1)**. A programmable magnetic angle sensor (Digikey) was attached for continuous monitoring of wheel motion. Headposts were custom-designed to mimic the natural head angle of the running mouse, and mice were mounted with the center of the body at the apex of the wheel. On each training day, a headpost was attached to the implanted nuts with two screws (McMaster-Carr). The headpost was then secured with thumb screws at two points on the wheel (**Fig. S1**). Mice were headposted in place for increasing intervals on each successive day. If signs of anxiety or distress were noted, the mouse was removed from the headpost and the training interval was not lengthened on the next day. Mice were trained on the wheel for up to 7 days or until they exhibited robust bouts of running activity during each session. Mice that continued to exhibit signs of distress were not used for awake electrophysiology sessions.

### Electrophysiology

All extracellular single-unit, multi-unit, and LFP recordings were made with an array of independently moveable tetrodes mounted in an Eckhorn Microdrive (Thomas Recording). Signals were digitized and recorded by a Digital Lynx system (Neuralynx). All data were sampled at 40kHz. All LFP recordings were referenced to the surface of the cortex. LFP data were recorded with open filters and single unit data was filtered from 600-9000Hz. Mouse pupil dilation was imaged in IR using a Flea 3.0 camera (Point Grey) with a zoom lens (M12 Lenses, Inc.). Illumination was generated by a positionable IR LED array (Mouser Electronics).

Awake recordings were made from mice that had received handling and wheel training as described above. On the initial recording day, a small craniotomy was made over V1 under light isoflurane anesthesia. The craniotomy was then covered with a cap, after which the mouse was allowed to recover for 2 hours. Mice were then fitted with a headpost and secured in place on the wheel apparatus (**Fig. S1**) before electrodes were lowered. Electrodes were initially lowered to ∼200 μm, then independently adjusted after a recovery period of 30-60 min. At the end of a recording session, the craniotomy was flushed with saline and capped. On subsequent recording days, the craniotomy was flushed with saline before placing the electrode array in a new site. Recordings were performed mainly in the second half of the light portion of the day/light cycle.

Visual stimuli were presented on an LCD monitor at a spatial resolution of 1680x1050, a real-time frame rate of 60Hz, and a mean luminance of 45 cd/m^2^ positioned 15cm from the eye. Stimuli were generated by custom-written software (J. Cardin, Matlab). Initial hand-mapping was performed to localize the receptive fields of identified cells. To maximize data collection, visual stimuli were positioned to cover as many identified receptive fields as possible. All stimuli were sinusoidal drifting gratings at a temporal frequency of 2 Hz, presented at a fixed interval of 1.5 s with an interstimulus interval of 2 s. We used either blocks of visual stimuli where orientation was held constant and contrast was varied, or where contrast was held at 100% and orientation was varied. Contrast tuned stimuli (5-20 levels) were optimized for mean orientation selectivity and spatial frequency. In a subset of experiments, a single orientated grating was repeatedly presented at 100% contrast.

For experiments in which air puffs were used to arouse the mouse, a small tube was positioned behind the head. Air puffs were delivered to the body of the mouse with minimum waiting intervals of 1-2 min with pseudorandom timing and jitter of 60 s. Air puffs were generated by a small oxygen tank using a solenoid-operated piston pump (Clark Solutions) to control timing and amplitude. The intensity of the air puff was adjusted during each experiment such that a noticeable increase in arousal – as indexed by pupil diameter – could be observed, without resulting in locomotion. For the air puff analyses, we only included trials that occurred during quiescence epochs, and had a maximum of 2 cm movement across the analyzed interval.

### Spike sorting

Spikes were clustered semi-automatically using the following procedure. We first used the KlustaKwik 3.0 software^59^ to identify a maximum of 30 clusters using the waveform Energy and Energy of the waveform’s first derivative as clustering features. We then used a modified version of the M-Clust environment to manually separate units. Units were accepted if a clear separation of the cell relative to all the other noise clusters was observed, which generally was the case when a conventional metric of cluster separation, isolation distance (ID)^60^ exceeded 20. We further ensured that maximum contamination of the ISI (Inter-spike-interval) histogram <1.5 ms was smaller than 0.1%. In a small number of cases we accepted clusters with isolation distances smaller than 20, which could be caused by e.g. non-Gaussian clusters, only if separation was of sufficient quality as judged by comparing the cluster with all other noise clusters (20%, 80% quantiles of ID = 19, 46; median±standard error of median = 25±1.1).

### Data analysis

All data was analyzed using the Mathworks Fieldtrip toolbox and custom-made C and Matlab scripts (M. Vinck).

#### Computation of wheel position and change points

Wheel position was extracted from the output of the angle sensor. Since wheel position is a circular variable, we first transformed the sensor data to the [-π,π] interval. Because the position data would make sudden jumps around values of π and – π, we further performed circular unwrapping of the position phases to create a linear variable.

We then used a change-point detection algorithm that detected statistical differences in the distribution of locomotion velocities across time. The motivation of this method relative to the standard method of using an arbitrary threshold (e.g., 1 cm/s;^15^) is that our technique allowed for small perturbations in locomotion speed to be identified that might otherwise fail to reach the locomotion threshold. Further, it ensured that the onset of locomotion could be detected before the speed reached 1 cm/s. If the distributions of data points 100 ms before and 100 ms after a certain time point *t* were significantly different from each other, using a standard T-test at p<0.05 and sampling at 2 kHz, then the data point was deemed a candidate change point. A point *t* was considered a candidate locomotion onset point if the speed 100 ms after *t* was significantly higher than 100 ms before *t*. A point was considered a candidate locomotion offset point if the speed 100 ms after *t* was significantly lower than 100 ms before *t*. A point was accepted as a locomotion onset point if the previous transition point was a locomotion offset point. A point was considered as a locomotion offset if it was preceded by a locomotion onset point, and if the speed 100 ms after *t* did not significantly differ from zero. This prevented a decrease in speed to be identified as a locomotion offset point. We further required that a locomotion offset point not be followed by a locomotion onset point for at least 2 s, because mice sometimes showed brief interruptions between bouts of running.

We selected locomotion trials for which the average speed until the next locomotion offset point exceeded 1 cm/s and which lasted longer than 2 s. Quiescence trials were selected that lasted longer than 5 s, had an average speed <1 cm/s, and for which the maximum range of movement was <3 cm across the complete quiescence trial.

#### Pupil diameter extraction

Pupil diameter was extracted from gray-scale video frames of 800 × 600 pixels at 10 fps. 1) A threshold on the luminance of 80 [scale 0 to 255] was set removing background pixels outside the eye. 2) After thresholding, fuzzy c-mean clustering was used to identify two clusters of pixels, which typically yielded one set of relatively dark pixels for the pupil and a set of relatively light pixels for the rest of the eye. In some cases, the shadows at the edges of the eye also fell in the set of dark pixels. 3) We then determined the two darkest spatial clusters of adjoined pixels (sum of inverted luminances on scale of 0 to 255). 4) In some sessions, an eyelash would be in the middle of the eye. In those cases, we used dilation (using MATLAB’s *imdilate* function) with a ball of 30 pixels that ensured that the regions would be adjoined. 5) Another problem was that the light caused a cluster of pixels with very high luminance around the pupil, which would create artificial edges and distort the shape of the pupil. We therefore allowed no edges to be detected within 20 pixels of other pixels having a luminance above 200. 6) Because the edges of the eye could be quite dark due to shadows, the darkest region could sometimes fall on the edge of the eye. Automatically selecting the region that was the darkest, had the highest contrast, or was the roundest (or a combined function of the three features) would give a good but not fully reliable identification of the pupil as judged by human observers. To solve this problem, we *post-hoc* set a region of interest constraining the location in which the pupil center was to be found. 7) We then fit an ellipse to the detected edges using least-squares fitting and determined pupil diameter as the surface of the ellipse. This procedure eliminated issues arising from missing data around the spot of high luminance caused by lighting (**Fig. 1 and 5**). 9) All video readouts were inspected for quality at 32x video speed, and few videos or segments of videos with poor readout quality (e.g. due to the mouse not fully opening its eyes) were rejected.

#### LFP power

For the analysis of time-courses around locomotion onset and offset, we estimated the LFP power spectral density at each time point. We first computed spectrograms using 7 cycles of LFP data per frequency and a Hann taper. We then performed smoothing of the spectra with rectangular box car windows such that always 4 s of data was used to estimate power at a certain time point. For example, for a frequency of 4 Hz, smoothing with 4 – 1.75 = 2.25 s was performed, while for a frequency of 50 Hz, 4 – 0.14 = 3.86 s smoothing was used. This procedure is comparable to Welch averaging or multitapering and ensures that time courses across frequencies can be compared, while preventing large variance in spectral estimation due to long estimation windows for high frequencies. For the locomotion onset and offset analysis in **Fig. 2** we only used sessions for which data was available for the full analyzed periods, i.e. 20 and 40 s after L-on and L-off, respectively. For computing LFP spectra during visual stimulation, we divided the data in 500 ms segments and used multitapering with ±4 Hz smoothing.

#### Noise correlations

Noise correlations were computed as in ^36^. For each unique visual stimulus, we Z-scored the firing rates, computed for the entire stimulus period, across presentations of that same stimulus. Only stimuli for which there were at least 3 presentations were considered. We then concatenated the Z-scored firing rates across the different unique stimuli. Noise correlation was then defined as the Pearson’s correlation coefficient. Note that using Spearman rank correlations gave qualitatively similar results.

#### Computation of modulation and SNR

Computation of firing rate modulation and SNR were always computed as:

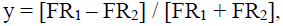

where FR1 and FR2 are the firing rates in two separate conditions (e.g. locomotion and quiescence, or visual stimulation and the ITI). The modulation and SNR are quantities that ranges between −1 and +1.

#### Quantification of burstiness

We quantified the propensity to engage in burst firing using the coefficient of Local Variation (LV^61^). The LV quantifies the irregularity of ISIs. A Poisson-process has an LV of 1. Rhythmic firing leads to LV values <1, whereas burst firing leads to LV values above 1^61^. The LV quantifies irregularity only on the basis of pairs of subsequent ISIs, and is therefore robust against non-stationarities in firing rates. We also computed an alternative measure of burstiness, as the log fraction of ISIs between 2-10 ms over the fraction of ISIs between 10 and 100 ms, Log(ISI_short_ / ISI_long_). We found a strong correlation between the latter measure of bursting and the LV for RS cells (Pearson’s R = 0.84, p=7×10^−15^, **Fig. S9**). Thus, bursting cells were characterized both by firing many spikes with short ISIs, and firing irregularly.

#### Computation of spike densities

Instantaneous firing rate was computed by convolving the spike trains either with a rectangular kernel or a Gaussian smoothing kernel. For tracking longer-lasting changes in firing rate around state transition points (**Fig. 1, 3, 7**), we used relatively long smoothing kernels (±500 ms Gaussian kernels, with SD of 250 ms). For visualizing neuronal responses to visual stimuli (**Fig. 4, 8**), we used short smoothing kernels (±25ms Gaussian kernels, with SD of 12.5ms). For computing cross-correlation coefficients between locomotion speed and firing rate, we used ±500 ms sliding box car windows to compute both speed and instantaneous firing rate.

#### Correlation of time, firing rate and power

We computed Spearman’s rho correlations between time after locomotion offset (>3 s) and instantaneous firing rate by computing the instantaneous firing rate every 500 ms with a Gaussian smoothing kernel of ±500 ms (SD of 250 ms). Taking steps of 500 ms ensured that datapoints were not correlated because of smoothing, which would have spuriously increased correlation coefficients. For the correlation with power and pupil diameter over time, we used steps of 4 s, since a ±2 s window was used to compute instantaneous power.

#### Statistical testing

Non-parametric statistics were used throughout the manuscript to avoid making assumptions inherent to parametrical testing, except for linear regression analysis, as we were explicitly interested in testing for linear relationships.

